# Spatial variation in population-density, movement and detectability of snow leopards in a multiple use landscape in Spiti Valley, Trans-Himalaya

**DOI:** 10.1101/2020.09.09.289181

**Authors:** Rishi Kumar Sharma, Koustubh Sharma, David Borchers, Yash Veer Bhatnagar, Kulbhushan Singh Suryawanshi, Charudutt Mishra

## Abstract

The endangered snow leopard *Panthera uncia* occurs in human use landscapes in the mountains of South and Central Asia. Conservationists generally agree that snow leopards must be conserved through a land-sharing approach, rather than land-sparing in the form of strictly protected areas. Effective conservation through land-sharing requires a good understanding of how snow leopards respond to human use of the landscape. Snow leopard density is expected to show spatial variation within a landscape because of variation in the intensity of human use and the quality of habitat. However, snow leopards have been difficult to enumerate and monitor. Variation in the density of snow leopards remains undocumented, and the impact of human use on their populations is poorly understood. We examined spatial variation in snow leopard density in Spiti Valley, an important snow leopard landscape in India, via spatially explicit capture recapture analysis of camera trap data. We camera trapped an area encompassing a minimum convex polygon of 953 km^2^. We estimated an overall density of 0.49 (95% CI: 0.39-0.73) adult snow leopards per 100 km^2^. Using AIC, our best model showed the density of snow leopards to depend on wild prey density, movement about activity centres to depend on altitude, and the expected number of encounters at the activity centre to depend on topography. Models that also used livestock biomass as a density covariate ranked second, but the effect of livestock was weak. Our results highlight the importance of maintaining high density pockets of wild prey populations in multiple use landscapes to enhance snow leopard conservation.

## Introduction

Large carnivores typically range over large areas (1), occur naturally at low densities (2) and exhibit elusive behavior. Approximately 60% of the world’s largest carnivores are threatened with extinction (3). Many large carnivore populations have undergone severe declines in their population size as well as distribution in the past few decades resulting in significant trophic cascades (4).

Evaluating the status of large carnivore species and the effectiveness of conservation actions requires rigorous monitoring of their populations. Inaccurate and imprecise estimates of population abundance can have larger cascading effects on conservation of endangered species by their potential to influence a range of scientific inferences as well as conservation interventions. However, large carnivores in general are difficult to enumerate due to their large ranges (1), naturally low densities (2), and elusive behaviour.

The threatened snow leopard *Panthera uncia* is a typical example of a difficult to sample, elusive carnivore that is reported to occur at relatively low population densities (0.15-3.88/100 km^2^) even in best habitats (5–7). Snow leopards have relatively large home ranges, and of the 170 protected areas in the global snow leopard range, 40% are smaller than the home range size of a single adult male (8). The distribution range of the snow leopard across Asia is subject to pervasive human use, predominantly in the form of pastoralism and agro-pastoralism (9). Over the past two decades, snow leopard habitats have also come under the increasing purview of developmental activities and mining (6), commercial livestock rearing such as cashmere goats (10), extraction of *Cordyceps*, and tourism.

Snow leopard habitats represent multiple use landscapes dominated by pastoralism and agro-pastoralism. Conservationists generally agree that snow leopards must be conserved amidst people, following a land-sharing approach, rather than too much emphasis on creating strictly protected areas (8). Such an approach, however, requires a good understanding of the impact of land use on snow leopard populations.

Within a landscape, snow leopard density is expected to show spatial variation because of variation in the intensity of human use and the quality of habitat. A good understanding of such variation and its correlates is important for designing appropriate, spatially explicit strategies for land-sharing. However, snow leopard population abundances have been difficult to estimate, and spatial variation in their density remains undocumented and poorly understood. In this study, we assessed the spatial variation in snow leopard density and examined its ecological correlates in Spiti Valley, one of India’s most important snow leopard landscapes. We used the Akaike Information Criterion (AIC) to select spatial capture-recapture (SCR) models that help explain spatial variation in density, encounter rate and habitat use. Our findings suggest that maintaining pockets of high density wild prey populations can immensely facilitate snow leopard conservation in multiple use landscapes.

## Materials and Methods

### Study area

Spiti Valley (31° 35’-33° 0’N; 77° 37’-78°35’E) is in the Indian state of Himachal Pradesh. All necessary research permits were received before conducting the field work from the Chief Wildlife Warden, Government of Himachal Pradesh, India. Comprising of approximately 12,000 km^2^ of catchment of the river Spiti. It is flanked by the Greater Himalaya in the south, Ladakh in the north and Tibet in the east. Lying in the rain-shadow of the Greater Himalaya, the region is cold and arid, with most of the precipitation in the form of snow. The main vegetation type is dry alpine steppe and the region is characterized by the absence of trees. The landscape is rugged and altitude ranges between 3000 to 6000 meters. Spiti experiences cold winters with temperature dropping below −30°C, while summers have a mean maximum temperature of about 25°C.

In our study area (Fig. 1), there were 50 hamlets and villages, with number of houses ranging from 2 to 231 and their human populations ranging from 7 to 706. The human population density in the valley is about one person per square kilometer. The local people are mainly agro-pastoralist, while parts of the valley in summers are used by transhumant pastoralists. Livestock species includes yak *Bos grunniens*, dzo (hybrid of cow and yak), dzomo (female dzo), cow *Bos indicus*, horse *Equus caballus*, goat *Carpa hircus*, sheep *Ovis aries* and donkey *E. asinus*. The main livestock grazing areas are located between 3,800 to 5,000 meters and communities have traditional grazing rights over rangelands. Wildlife of the region includes wolf *Canis lupus*, ibex *Carpa sibirica*, bharal *Psedois nayaur*, hare *Lepus oiostolus*, and golden eagle *Aquila chrysaetos*.

**Figure 1.**
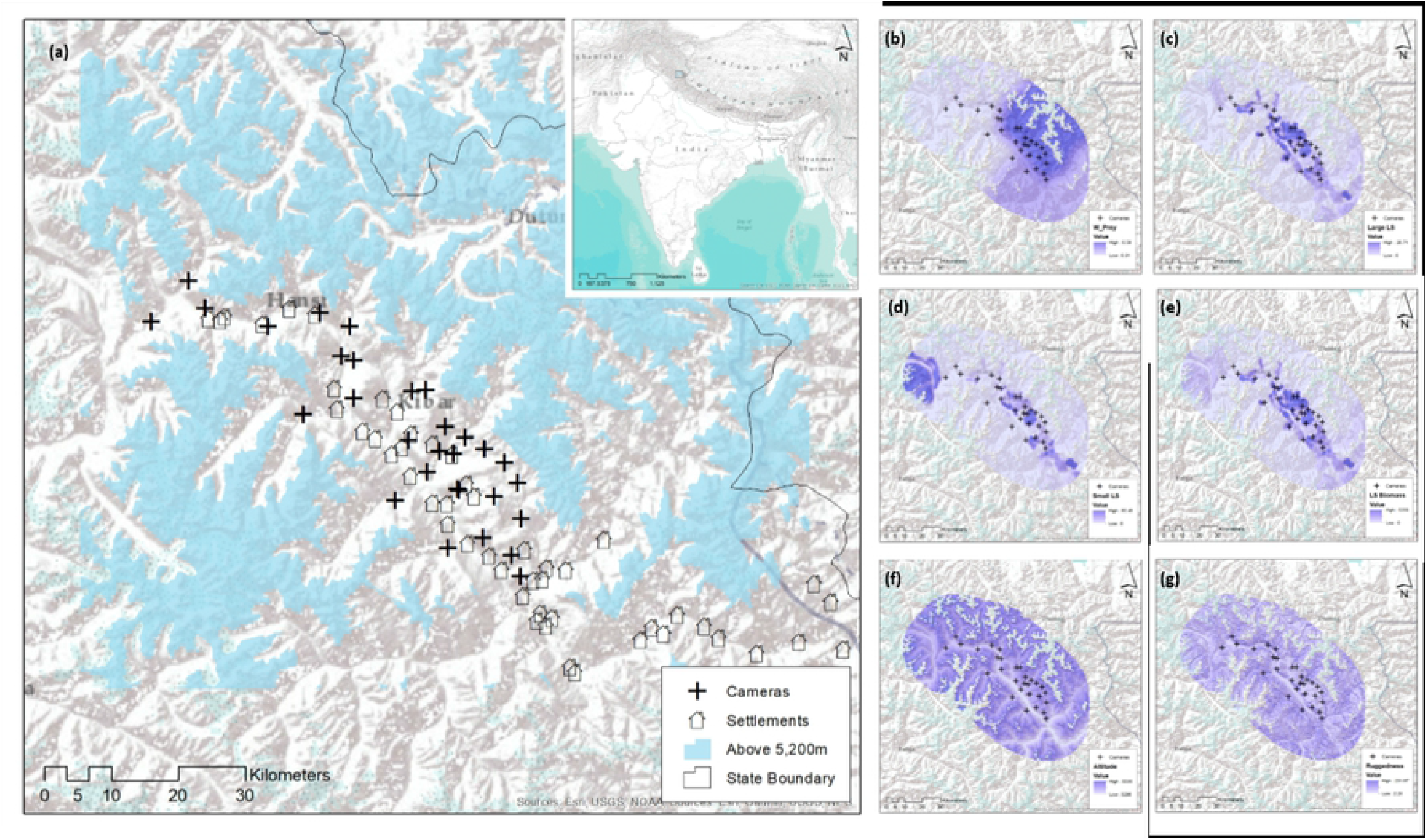
(a) Map of the study area showing camera trap locations and sampling region characterized by areas below 5200 meters. The inset map shows location of the study area in the state of Himachal Pradesh, India, and maps (b), (c), (d), (e), (f) and (g) show spatial variation in wild prey density, density of large bodied livestock, density of small bodied livestock, overall livestock biomass, altitude and terrain ruggedness respectively within the area of integration.

Snow leopards and wolves were historically persecuted in the region in retaliation for livestock depredation, though retaliatory killing has declined substantially owing to community-based conservation efforts.

### Estimating snow leopard population density using camera traps

We deployed Reconyx RM45 camera traps at 30 locations over an area of 953 km^2^ (Minimum Convex Polygon joining the outermost trap locations) with an average inter-trap distance of 4035 m (SE = 374m) (Fig. 1). The camera traps were deployed from October 2011 to January 2012 for a period of 80 days with an overall trap density of 3 camera traps per 100 km^2^ following recommendations of placing at least two traps per average home range (11) or at least 2 traps per average female home range (12). The cameras traps were deployed at sites where we encountered relatively high frequency of snow leopard signs such as scrapes, pugmarks, scats and scent marks, especially around terrain features that snow leopards are known to prefer for marking and movement such as ridgelines, cliffs and gully beds. We used a combination of single side (n=14) and both side (n=16) camera trap placement to optimize coverage and identification of individuals. Our cameras recorded snow leopards at 25 out of 30 locations without using any baits or scent lures.

Individual snow leopards captured in the images were identified based on their pelage patterns by two independent observers using at least three similarities or differences (5,13). A total of six encounters were discarded as the images were not good enough to identify the individuals. Following concerns raised by Johansson et al. (2020), we used the *Snow Leopard Identification: Training and Evaluation Toolkit* (https://camtraining.globalsnowleopard.org/leppe/login/) to test the skills of both of our observers in identifying snow leopards. Our observers scored 96.3% and 88.9% accuracy respectively in identifying snow leopards from 40 blind, independent trials, thus leaving us confident of identifying individuals with reasonable accuracy. Snow leopard capture histories were built using the standard count detector format of the secr package in R (Efford 2017) where each encounter of an identified cat was linked to a detector (camera trap), whose location, period of operation and other relevant covariates were recorded in a separate table. We restricted the study period to 80 days and assumed that the population was closed and that there was no temporal effect on detection probability of snow leopards during the sampling period.

Typically, SCR models assume that expected encounter rate depends on the Euclidean distance between detector and activity centre, but in highly structured environments such as steep mountains, this may not always be true. For example, we may record more encounters for a snow leopard in a distant trap than a closer trap if the habitat between the closer trap and activity centre has greater resistance to movement (e.g. a deep gorge separating two detector locations). Royle et al. (2013) and Sutherland et al. (2015) proposed replacing Euclidian distance with a least-cost path distance (ecological distance) in which movement cost depends on the habitat. The method involves estimation of movement cost parameter(s) simultaneously with other SCR parameters. Sutherland et al. (2015) demonstrated that violations of the Euclidean distance assumption can bias estimates of density and they suggest that least-cost distance be tested in highly structured landscapes.

We used the maximum likelihood based SCR models (14) to estimate density while investigating the effect of least-cost path distance on movement, using package ‘secr’ (15) in R (16). The method involves integration over a 2-dimensional region containing the possible (and unknown) locations of the activity centres of animals at risk of detection. The region of integration is based on a polygon extending a certain distance (the buffer width) beyond the outermost traps.

We used the inbuilt ‘suggest buffer’ function of secr to arrive at a buffer of 24,000 meters assuming it to be wide enough to keep any bias in estimated densities as acceptably small (i.e. snow leopards with activity centres beyond 24 km from the outermost traps had a negligible probability of being captured in the detectors). Areas above 5,200 m from mean sea level were treated as non-habitat because these areas are mostly devoid of vegetation and prey species. We defined an integration area with spacing of 500 m, assuming that snow leopard density was not likely to change at a finer resolution. We used the top model chosen by minimum AIC to estimate population size (*N*) and density (*D*) over the integration region (14).

### Spatial capture recapture models

The spatial distribution model in SCR is a spatial Poisson process for animal activity centres whose intensity (expected number of animal activity centres per unit area) can be homogeneous (constant over space) or inhomogeneous (varying over space) (Borchers and Efford 2008). We use the notation *D*(***x**; **θ***) for density, signifying that density is a function of activity centre location, ***x***, which is a vector representing the x and y coordinates of an activity centre, and of parameters represented by the vector ***θ***.

We fitted SCR models with various combinations of covariates defined a priori. A candidate model set was developed to investigate the effects of various covariates potentially influencing snow leopard behaviour, ecology and natural history. We investigated models with various combinations of covariates for the density model, the encounter function intercept model, and the encounter function range model. The general forms of the density model, and encounter function intercept and range models, respectively, are as follows:

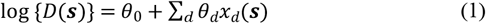

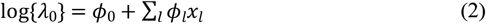

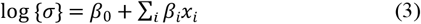

where

*x_d_*(*s*) is the *d*th spatially referenced covariate at location ***s*** that affects density (*D*), and *θ*_0_ and *θ_d_* are the density intercept parameter and *d*th regression parameter (all covariates were treated as known);
*x_l_* is the *l*th covariate that affects expected encounter rate at distance zero (*λ*_0_), and *ϕ*_0_ and *ϕ_l_* are the intercept parameter and *l*th regression parameter for expected encounter rate at distance zero;
*x_i_* is the *i*th covariate that affects the range parameter (*σ*), and *β*_0_ and *β_i_* are the range intercept parameter and *i*th regression parameter.

Half-normal encounter function forms were used, such that the expected number of encounters of an animal at a camera that is a distance *d* from its activity centre is *E*(*n*) = *λ*_0_exp { – *d*^2^(2*σ*^2^}.

For snow leopard density, we considered models in which the *x_d_*(***s***)*s* were wild prey density, livestock density, terrain ruggedness and altitude at ***s***. We investigated the effect of topographies (a factor with levels “ridgeline”, “cliff” or “gully bed”) and different altitudes at the activity centre on the encounter function intercept and range parameters. We also investigated models in which movement cost depends on altitude.

We modelled *D*(***s***) as a function of six spatial covariates (*x_d_*(***s***)*s*) that could affect snow leopard density (Figure 1). These included terrain ruggedness (typical snow leopard habitats are steep and rugged (18)), altitude (snow leopard densities are known to be a function of altitude (19)), wild prey density (believed to be the main determinant of snow leopard population abundance (20)), stocking density of large-bodied livestock and small-bodied livestock (potential prey for snow leopards, source of disturbance, and competitors for wild prey (18,21)).

Terrain ruggedness was derived using the terrain ruggedness index (22) from a 30 × 30 meter Digital Elevation Model from Aster Global Digital Elevation Model data using the terrain analysis plugin in the Quantum GIS 2.14 software (23). Livestock density was determined through door to door censuses in 51 villages in the study area. The pastures used by each village were mapped using Google Earth and livestock stocking densities for small- and large-bodied livestock were computed separately (as they are often herded separately (24)) by dividing the total livestock heads using a pasture, by the area of the pasture in square kilometres. We used average biomass of large-bodied and small-bodied livestock (25) to estimate the livestock biomass availability to snow leopards across the study area. Averaging over 1km, we smoothened the livestock density surfaces across the study area.

Abundance of wild prey, which primarily included blue sheep and ibex, was estimated using the double observer survey technique for the entire study area (26) (Table 1) between April and June 2012. Four teams, comprising two observers each, carried out the surveys for a period of eight days to cover the entire study area. Observers recorded the GPS coordinates of the sightings, the group size and age-sex classification of the groups encountered. The unique identity of each observed ungulate group was established through immediate post survey discussions between two observers using the age-sex classification, size and the location information of sightings (26). The study area was divided *post hoc* into 7 blocks delineated based on natural topographic features in the landscape. For each block, the cumulative number of wild ungulates encountered by the two observers were calculated. The relative density of wild ungulates for each block was estimated as total number of wild prey in a block divided by the size of the survey block (Table 2). The wild prey density surface was smoothened by averaging over a moving window of 5 km.

**Table 1.** Estimates of abundance of blue sheep and ibex using double observer approach in Spiti valley.

| Variable | Blue sheep | Ibex | Overall<br>(Blue sheep & Ibex) |
| --- | --- | --- | --- |
| $C$ | 75 | 18 | 93 |
| $S_1$ | 18 | 5 | 23 |
| $S_2$ | 6 | 1 | 7 |
| $\hat{G}$ | 100.42 | 24.26 | 124.71 |
| $Var(\hat{G})$ | 11.87 | 0.33 | 2.27 |
| $\hat{U}$ | 14.85 | 12.38 | 14.37 |
| $Var(\hat{U})$ | 1.55 | 2.19 | 1.09 |
| N (Naïve Estimate) | 1470 | 297 | 1767 |
| $\hat{N}$ | 1491 | 300 | 1792 |
| $Var(\hat{N})$ | 16067.87 | 1338.53 | 17482.66 |
| $\pm 95\%$ Confidence Interval | 251.64 | 72.63 | 262.49 |
| $P_1$ | 0.92 | 0.94 | 0.93 |
| $P_2$ | 0.80 | 0.78 | 0.80 |

$C$ , Number of groups seen in both surveys; $S_1$ , number of groups seen in first survey only; $S_2$ , number of groups seen in second survey only; $\hat{G}$ , estimated number of groups; N, Naïve population estimate; $\hat{N}$ , estimated population size; $P_1$ , $P_2$ , mean of the estimated detection probability for observer one and two, respectively.

**Table 2.** The seven regions of Spiti Valley showing variation in wild ungulate abundance and variation in snow leopard density.

| Region | Area<br>(km<sup>2</sup>) | Wild Prey<br>Density (per<br>km<sup>2</sup>) | LCL | UCL |
| --- | --- | --- | --- | --- |
| Chandertal | 768 | 0.01 | 0.01 | 0.01 |
| Kibber Plateau | 1623 | 0.58 | 0.49 | 0.66 |
| Dhar Ula | 578 | 0.29 | 0.25 | 0.34 |
|  |  | 0.23 | 0.19 | 0.26 |
| Dhankar<br>Lalung | 513 |  |  |  |
| Dhar Pangmo | 1423 | 0.01 | 0.01 | 0.01 |
| Guiling | 346 | 0.25 | 0.21 | 0.28 |
| Lossar Kiato | 1117 | 0.04 | 0.03 | 0.04 |

LCL= 95% lower confidence Limit
UCL=95% upper confidence limit

We developed an *a priori* set of models that we anticipated would best explain the variation in the density of snow leopards. Our global (most complex) model included terrain ruggedness, density of wild prey, stocking density of small-bodied livestock, stocking density of large-bodied livestock, and cumulative livestock biomass. We then fitted 15 candidate sub-models using subsets of the variables used in the global model. Each candidate sub-model represented a specific hypothesis about the relationship between snow leopard density and how snow leopards use space about their activity centres, and explanatory variables. We used Akaike’s Information Criterion (AIC) for model selection (27). All data analysis was implemented using package secr (15) in program R (16)

## Results

The double observer surveys for wild prey yielded abundances of 300±72.63 (95% CI) ibex and 1491± 251.64 blue sheep in the entire study area (Table 1). The naïve wild prey densities within the survey blocks ranged from 0.01 to 0.58 per km^2^ (Table 2). The terrain ruggedness across the integration region ranged from 5.39 to 120.83 (Mean= 37.62, SD = 18.81), the distance from nearest village from 251 meters to 3200 meters (Mean= 12241, SD = 8439). The density of large livestock ranged from 0 to 14.11 per km^2^ (Mean= 1.41, SD = 2.21) while that of the small bodied livestock ranged between 0 to 11.39 (Mean= 0.89, SD = 1.75) per km^2^. The mean livestock biomass ranged between 0 and 5,539 kg (Mean = 230.71, SD = 471.06).

We obtained 112 captures of 16 adult snow leopards over a sampling period of 80 days. Snow leopards were captured at 25 of the 30 camera trap locations. The estimate of snow leopard abundance was 25 (95% CI: 20-38) for an area of 5,144 km^2^ covering the area of integration. Our camera traps spanned the covariate space of the wild prey density reasonably well (fig. 2).

**Figure 2.**
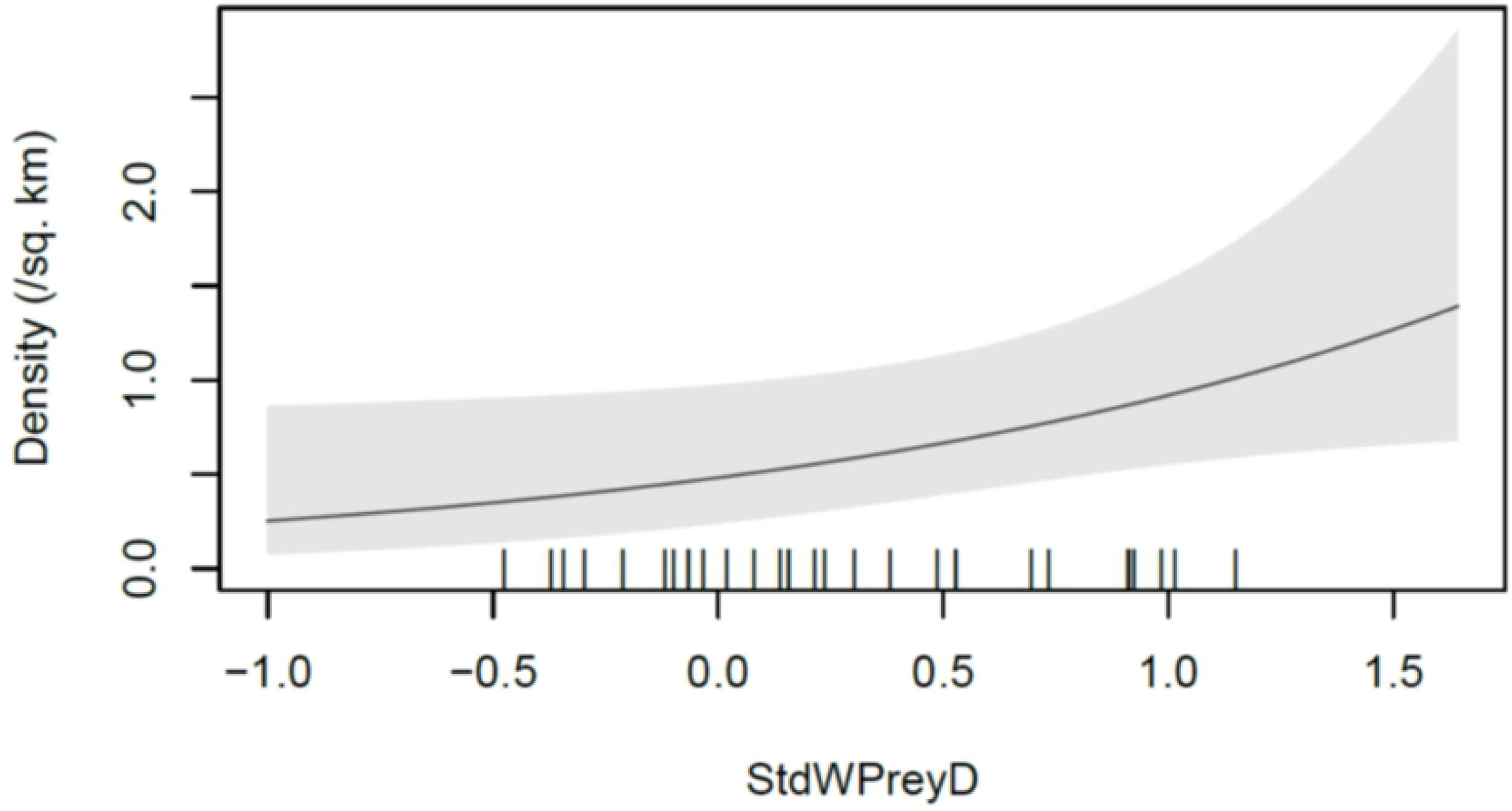
Snow leopard density estimated as a function of wild prey density (StdWPreyD) in Spiti Valley, India. The tick-marks on the x-axis show the placement of camera traps in the StdWPreyD dimension, with the range of the StdWPreyD axis indicating the range of StdWPreyD values in the data.

The density estimates from the top model ranged from 0.23 to 1.08 per 100 km^2^ across the region of integration (Table 2). The average snow leopard density for the study area was estimated to be 0.49 (SE=0.08) per 100 km^2^. The top 7 models with a cumulative AIC weight of 100% predicted snow leopard habitat use around their activity centres to be a function of altitude: The conductance coefficient associated with ecological distance in the best model (parameter *α*_2_ of Royle et al. (2013) and Sutherland et al., (2015) was estimated to be 0.38 (SE = 0.10), indicating that relatively higher altitudes within the study area boundaries were more conducive to snow leopard movement. Similarly, all top models used topography as a covariate affecting encounter rate at an activity centre. The models using wild prey density as a covariate affecting snow leopard density had a cumulative AIC weight of 0.78, followed by livestock biomass (AIC weight = 0.36) and then other covariates (Table 3). The coefficients for livestock biomass, however, were not significant at the 5% level. The top model, with AIC weight 0.45, indicated that wild prey density positively affected snow leopard density across the landscape (estimated *θ_prey_* = 0.59, SE = 0.27). On cliffs, the expected encounter rate for a camera at the activity centre was 0.083 (95% CI: 0.05-0.15), while it was 0.16 (95% CI: 0.11-0.23) in gully beds, and 0.22 (95% CI: 0.15, 0.32) on ridgelines.

**Table 3.** Models ranked on the basis of Akaike’s Information Criterion (AIC) for spatially explicit capture recapture estimates of snow leopard density in a multiple use landscape. Spatial capture recapture models are described using the following notation: “~1” shows the RHS of Equations (1) to (3) to only contain an intercept term; “~x” means that it contains an intercept and covariate “x”; “~x+y” means that it contains an intercept and covariates “x” and “y”; “x*y” indicates an interaction between x and y, npar = number of parameters in the model; and logLik = the model’s log-likelihood. The difference between the AIC and the minimum AIC for the given candidate model set is denoted by dAIC, while the associated weight is denoted by AICwt. D~, lambda0~, sigma~ and noneuc~ represent the density model, the encounter function intercept model, the encounter function range model and the conductance model respectively, modelled as functions of covariates or only a constant term.

| Model | npar | logLik | AIC | AICc | dAIC | AICwt |
| --- | --- | --- | --- | --- | --- | --- |
| D~WildPrey, lambda0~Topo, sigma~1, noneuc~Altitude | 7 | -154.1 | 322.18 | 336.18 | 0 | 0.45 |
| D~WildPrey + LSBiomass, lambda0~Topo, sigma~1, noneuc~Altitude | 8 | -153.7 | 323.5 | 344.1 | 1.30 | 0.24 |
| D~1, lambda0~Topo, sigma~1, noneuc~Altitude | 6 | -156.6 | 325.3 | 334.6 | 3.07 | 0.10 |
| D~WildPrey * LSBiomass, lambda0~Topo, sigma~1, noneuc~Altitude | 9 | -153.7 | 325.5 | 355.5 | 3.30 | 0.09 |
| D~SmallLS, lambda0~Topo, sigma~1, noneuc~Altitude | 7 | -156.2 | 326.4 | 340.4 | 4.25 | 0.05 |
| D~LargeLS, lambda0~Topo, sigma~1, noneuc~Altitude | 7 | -156.6 | 327.2 | 341.2 | 5.06 | 0.04 |
| D~LSBiomass, lambda0~Topo, sigma~1, noneuc~Altitude | 7 | -156.6 | 327.3 | 341.3 | 5.06 | 0.04 |
| D~WildPrey, lambda0~Topo, sigma~1 | 6 | -160.5 | 333.0 | 342.3 | 10.78 | 0 |
| D~Altitude + WildPrey, lambda0~Topo, sigma~1 | 7 | -159.9 | 333.8 | 347.8 | 11.60 | 0 |
| D~1, lambda0~Topo, sigma~1 | 5 | -162.7 | 335.3 | 341.3 | 13.12 | 0 |
| D~Altitude + WildPrey + Ruggedness, lambda0~Topo, sigma~1 | 8 | -159.8 | 335.7 | 356.2 | 13.46 | 0 |
| D~Altitude, lambda0~Topo, sigma~1 | 6 | -162.3 | 336.6 | 345.9 | 14.41 | 0 |
| D~Ruggedness, lambda0~Topo, sigma~1 | 6 | -162.3 | 336.7 | 346.0 | 14.47 | 0 |
| D~Altitude + Altitude<sup>2</sup>, lambda0~Topo, sigma~1 | 7 | -161.8 | 337.7 | 351.7 | 15.48 | 0 |
| D~1, lambda0~1, sigma~1 | 3 | -250.5 | 507.0 | 509.0 | 184.78 | 0 |

Ruggedness = Terrain Ruggedness Index, Altitude = Elevation above mean sea level, WildPrey = Density of wild prey, LargeLS = Stocking density of large bodies livestock, SmallLS = Stocking density of small bodied livestock, and LSBiomass = total livestock biomass; and Topo is the trap covariate depicting the placement of a camera trap at a ridgeline, cliff or gully bed.

Only 18% of the entire area of integration had estimated snow leopard density greater than 1 animal per 100 km^2^ from the top model, while 42% had estimated density lower than 0.25 animals per 100 km^2^.

## Discussion

Our study established the first baseline estimate of the population and density of the snow leopard in Spiti Valley, an important snow leopard habitat in India that has been identified by the Indian Government as a priority landscape under the Global Snow Leopard and Ecosystem Protection Program (28). In our study area, a combination of community-based conservation efforts over the years, peoples’ religious beliefs, and law enforcement have led to a near cessation of retaliatory killing of snow leopards and hunting of ungulates (Mishra 2016). The estimated snow leopard density in our study area was lower than that from studies conducted in several other smaller study areas (13,30–32), but there was considerable spatial variation in density in our study area. Our results support the possibility that density estimates from several earlier studies might be positively biased because of small study areas (< 400 km^2^) (13,33,34) that were located in high density parts of respective landscapes (Suryawanshi et al. 2019).

All the top models in our study indicated that conductance is greater at higher altitudes. Ecologically, this can be translated as snow leopards tending to move greater distances at higher altitudes, which matches natural history observations that suggest snow leopards to move along ridgelines (36–38).

Our top model showed that the variation in snow leopard density was largely associated with variation in wild prey density. It appears therefore that in multiple-use areas where killing of snow leopards is not a serious threat, the variation in the abundance of wild prey is the main determinant of spatial variation in snow leopard density. Models that included livestock biomass availability in addition to wild prey density were a close second. The negative coefficient of livestock biomass availability suggests that snow leopard densities tend to be lower in areas with high livestock density, although this coefficient was not significant at the 5% level. Other variables did not have any noteworthy effect on the snow leopard density within the study area. This is broadly in line with the conclusions of Suryawanshi et al. (2017), who have shown that snow leopard abundance is primarily determined by the abundance of wild prey, and not by the abundance of livestock. The models also indicate the possibility that livestock at high density may negatively influence snow leopard abundance through forage competition with wild ungulates (Mishra et al. 2004), or potentially high risk of conflict with humans leading to mortality. Snow leopard activity as well as wild prey densities are reported to be lower in areas with high livestock density (39).

Although snow leopards are known to prefer rugged terrain (40–42) we did not find much support for snow leopard density to be dependent on ruggedness. This is presumably because our entire study area was reasonably high on typical scales of ruggedness estimates.

Human settlements and associated anthropogenic pressures are considered to have a negative influence on carnivore habitat use (43,44). In the case of snow leopards, studies report conflicting results. For instance while one study found human settlements to exert a negative influence on snow leopard habitat use (45), other studies reported no such effect (46,47). In our study area, human density was low (<2 per km^2^), and livestock grazing the major anthropogenic activity. To some extent, the density of snow leopard activity centres in our study was marginally higher in areas with low livestock biomass.

Mishra et al. (2009) provided a conceptual framework for a land-sharing approach for wildlife conservation in snow leopard landscapes, that advocates maintaining a matrix of ‘core’ (no grazing or human use) and ‘buffer’ landscape units (grazing and other sustainable human use activities) maintained with community support. Our results suggest that this would be particularly useful in the south-east and north-west parts of Spiti Valley that have low snow leopard density (Fig. 3). There is evidence that creation of such ‘core’ landscape units with community support can lead to the recovery of wild prey, and therefore, of snow leopards (49). Such efforts require building long term partnerships with local communities by co-opting them in conservation efforts (50).

**Figure 3.**
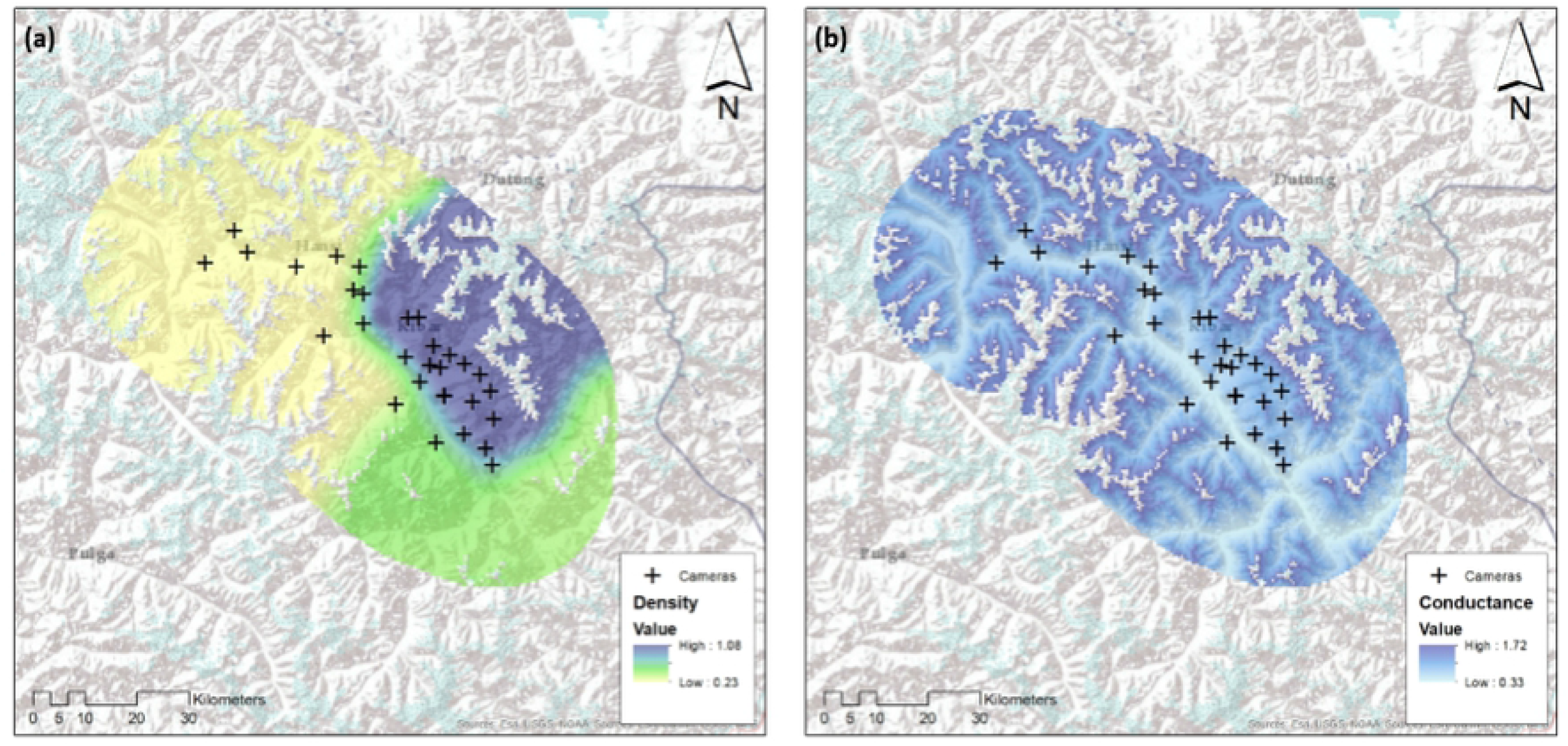
Maps of estimated density and conductance in the Spiti valley based on the top SCR model. (a) Snow leopard density, (b) log of conductance in the habitat for snow leopard movement. The coloured region shows the area of integration.

We suggest that the land sharing approach to snow leopard conservation can be strengthened considerably in snow leopard landscapes of Asia by creating core landscape units that can facilitate the recovery of ungulate populations.

## Acknowledgement

Whitley-Fund for Nature, Panthera and Snow Leopard Network provided primary support to this project. We are thankful to the Chief Wildlife Warden, Himachal, Divisional Forest Officer, Kaza and the Range Officer, Kaza, for permissions and logistics. Chunnit Kesang, Tenzin Thukten, Rinchen Tobgey, Sushil Dorje, Chudim and Takpa provided tremendous support in field.

